# RintC: fast and accuracy-aware decomposition of distributions of RNA secondary structures with extended logsumexp

**DOI:** 10.1101/656256

**Authors:** Hiroki Takizawa, Junichi Iwakiri, Kiyoshi Asai

## Abstract

The analysis of secondary structures is essential to understanding the function of RNAs. Because RNA molecules thermally fluctuate, it is necessary to analyze the probability distribution of secondary structures. Existing methods, however, are not applicable to long RNAs owing to their high computational complexity. Additionally, previous research has suffered from two numerical difficulties: overflow and significant numerical error. In this research, we reduced the computational complexity in calculating the landscape of the probability distribution of secondary structures by introducing a maximum-span constraint. In addition, we resolved numerical computation problems through two techniques: extended logsumexp and accuracy-guaranteed numerical computation. We analyzed the stability of the secondary structures of 16S ribosomal RNAs at various temperatures without overflow. The results obtained are consistent with *in vivo* assay results reported in previous research. Furthermore, we quantitatively assessed numerical stability using our method. These results demonstrate that the proposed method is applicable to long RNAs. Source code is available on https://github.com/eukaryo/rintc.

## Introduction

Functional noncoding RNAs (ncRNAs) play essential roles in a wide range of biological phenomena. Secondary structures are often crucial for the function of RNAs. There are a number of studies and software tools that can predict a single secondary structure for a given RNA. According to detailed analyses of free energy, however, some RNAs do not always form a single stable structure. Therefore, quantitative evaluations of the fluctuation of RNA secondary structures have recently attracted attention. Recent studies have provided methods to analyze the distribution of RNA secondary structures in more detail using the marginal probability of the Hamming distance(1), (2), (3).

The computational costs of basic algorithms used by previous methods are high, but the Fourier transform has been shown to reduce the time complexity of analysis. (4), (5), (6) Those previous methods, however, cannot be used to analyze long ncRNAs (e.g., > 1000 nucleotides) because the computational complexities are still too high. In this paper, we show that further reduction of computational complexity is possible by introducing maximum-span constraint on base-pairs (7).

While implementing and experimenting with our proposed method, we encountered two numerical computation problems. The first problem consists of numerical errors caused by Fourier transform. Specifically, the magnitude of numerical errors is uniform across large and small marginal probabilities, resulting in the small marginal probabilities being unreliable. This phenomenon occurs because the Fourier transform distributes numerical errors in its dynamic programming post-processing. A previous research (8) mentioned this type of numerical instability, but they have not shown detailed analysis. Accuracy-guaranteed numerical computation may provide quantitative evaluation of numerical instability, but this was unexamined by previous studies. In the context of kinetic analyses, for example, meta-stable structures are particularly interesting (9). Meta-stable structures may have considerably high free energy compared to the minimum free energy structure. In such a case, the Boltzmann probability of the meta-stable region can be very small. For reliable evaluation of meta-stable regions using distance-wise decomposition like RintW, quantitative assessment of the numerical error introduced by the Fourier transform method is necessary.

Interval arithmetic is a method in which arithmetic operations are defined along intervals expressing numerical values between the upper/lower edges. The approximate calculation of pi by Archimedes in the 3rd century BC is known as the oldest example of interval arithmetic. Around the 1950s, interval arithmetic came to be used for estimating the upper bounds on the numerical error caused by floating-point arithmetic in computers. For example, Sunaga (10) published one of the first studies in English on comprehensive algorithms for interval arithmetic for computers. Numerical calculations, however, have been used in only limited research fields, and few early studies were published.

The other numerical problem is that overflow occurs for long sequences. In stochastic models such as hidden Markov models common throughout bioinformatics, logsumexp (logarithm of summation of exponentials) is the standard solution to prevent overflow or underflow in numerical calculation. There is a limitation, however, in that it cannot handle zero nor negative values. This limitation is a problem when processing complex numbers with rectangular coordinates in Fourier transform. One solution is to apply logsumexp only to radii using polar coordinates, but simple application of polar coordinates causes problems when combined with interval arithmetic for accuracy-guaranteed numerical computation. Complicated conditions occur when the angular interval crosses zero or the radius interval contains zero. In this paper, in addition to a radius of polar coordinates, normalized orthogonal coordinates, rather than angles, are combined. Consequently, we can realize the advantages of logsumexp and interval arithmetic while preserving the simplicity of implementation.

We introduced maximum-span constraint to RintW in order to reduce the computational costs. While Turner energy parameters have been determined by experiments using short RNA, the computational analysis tends to be unstable when RNA can form long range base-pairs (11). Our method lost little by excluding long range base-pairs.

## Methods

### A. RintW + maximum-span

At first, we introduced maximum-span constraint in base pairs to the baseline algorithm of RintW (6). Detailed descriptions of RintW and the proposed method are in the accompanying supplementary file. The inputs of the algorithm are an RNA sequence and a reference secondary structure, and the outputs are the existence probability and the base pairing probability matrix for each Hamming distance from the reference secondary structure.

#### A.1. the representation of RNA secondary structure

As a computer-efficient expression, the RNA secondary structure was represented by a binary upper triangular matrix *σ* where each element was {0, 1}. Each element of *σ* was decided as follows.

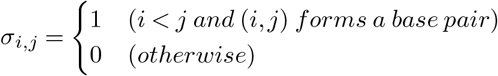

The distance between two RNA secondary structures *σ*_1_, *σ*_2_ were determined by the number of elements with different values, namely, Hamming distance.

It was also possible to use distance other than Hamming distance (5), but this time Hamming distance was used. The reason is that when the RNA secondary structure changes with time, it can be considered to be limited to the change of the Hamming distance 1, that is, whether one base pair is generated or extinguished. Hamming distance is a natural distance in that sense. Also, many previous studies (1), (4), (6) used Hamming distance, which was considered to be a standard method. For example, 5’-3’ distance (12) is known as a distance other than the Hamming distance, but introducing this was out of scope of this work.

Only secondary structures that satisfy the following constraints were considered. *N* = sequence length.

1. Watson-Crick base pairs(A-T, C-G) or wobble base pairs(G-U) only exist.
2. (prohibition of pseudo-knot) for all 1 ≤ *i* < *j* < *k* < *l* ≤ *N*, (*i*, *k*) and (*j*, *l*) do not form base pairs at the same time.
3. (max loop constraint) for all 1 ≤ *i* < *j* < *k* < *l* ≤ *N*, if (*i*, *l*) and (*j*, *k*) form base pairs and no paired base exists between *i* + 1 and *j* − 1 nor between *k* + 1 and *l* − 1, then *j* − *i* + *l* − *k* ≤ *C* + 2. *C* is a max-loop parameter, and We used *C* = 30 following many previous works.
4. (max span constraint) for all 1 ≤ *i* < *j* ≤ *N*, if (*i*, *j*) forms a base pair, then *j* − *i* ≤ *W*.

1. and 2. were the standard constraints used in the previous methods (1), (4), (6). 3. is called max loop constraint. This constraint was adopted by many of RNA secondary structure analysis methods using the energy model (13) described in the next section. This constraint reduces the time complexity. It is empirically known that this constraint has little effect on the calculation result. 4. is a constraint being studied in other previous works (14), (15), (7), (16), (11), but it was not used in the research by RintW(6) and we introduced it. This constraint is considered to be suitable for examining local structural motifs. (11))

#### A.2. energy model

Each structure was considered to have energy, and this Boltzmann probability was analyzed. An energy model (13) that can be analyzed by dynamic programming was adopted. In this model, the energy of the secondary structure was expressed by the sum of the following functions.

1. *f*_*h*_(*i*, *j*) = the energy of base pair (*i*, *j*) forming a hair-pin loop.
2. *f*_*l*_(*i*, *j*, *k*, *l*) = the energy of base pair (*i*, *l*) and (*j*, *k*) making a 2-Loop when *i* < *j* < *k* < *l*.
3. *f*_*mc*_ = the energy of having one multi loop.
4. *f*_*mi*_ = the energy of having one internal branch of multi loop.
5. *f*_*d*_(*i*, *j*) = the energy of base pair (*i*, *j*) forming a multi loop or being an outermost base pair.

#### A.3. polynomial approach

In previous researches (4), (5), (6), the polynomial approach was used as a method to reduce the time complexity of dynamic programming. For details of the approach, see the description of the previous articles. Briefly, a naïve dynamic programming method requires a convolution operation. This operation is regarded as computation in the spatial domain and is expressed by calculation in the frequency domain. Convolution operation can be converted to an inner product, and the computational complexity is reduced. After completing the dynamic programming computation, shift to the spatial domain by performing the Fourier transform. The same method was used in this study.

#### A.4. Preprocessing

As in the RintW algorithm, we calculate the following 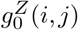 functions in *O*(*N*^2^) time as preprocessing, to obtain the gains of the Hamming distance 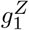 to 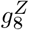 and 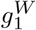 to 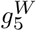 for (1 ≤ *i* ≤ *N*)

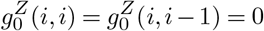

for (1 ≤ *i* < *j* ≤ *N*)

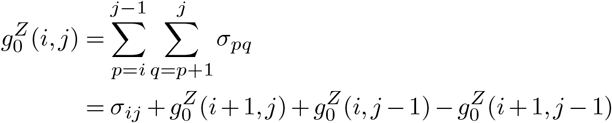

where *σ* is a binary matrix representation of the reference secondary structure. The maximum Hamming distance of the secondary structure from the representative secondary structure (*H*_*max*_) is also computed at this time (5).

#### A.5. Definitions of Function gs

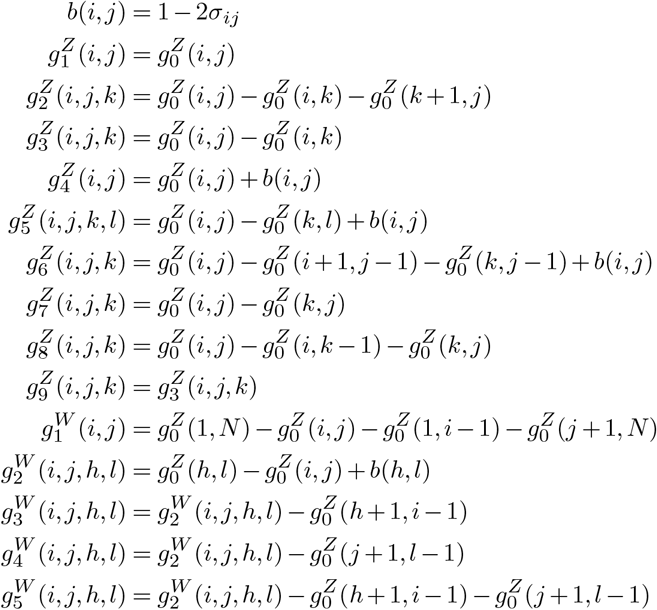

These functions calculate the Hamming distance of a substructure from the reference substructure. These functions were also used in previous researches. (5), (6) According to Mori *et al.* research (5), by changing this function, one can decompose the structures by another distance indicator. (*i.e.* other than the Hamming distance) This indicates a further potential of this concept, but it was out of the scope of this study to examine it.

#### A.6. Dynamic programming of the partition function

In the following equations, *x* is the (*H*_*max*_ + 1)-th root of unity. If Cooley–Tukey fast Fourier transform (FFT) is used instead of discrete Fourier transform (DFT) in post-processing, *x* is the smallest power of 2 that is equal to or greater than (*H*_*max*_ + 1). There are (*H*_*max*_ + 1) kinds of (*H*_*max*_ + 1)-th roots of unity calculated independently. Therefore, parallel computation is possible.

In order to avoid overflow, the proposed extended logsumexp is used. In the following equations, 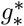 and 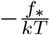 are real numbers, but 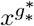 and 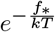 are converted into complex logsumexp type. Consequently, All DP-variable 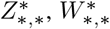, and 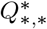 are also complex logsumexp type.

Initialization:

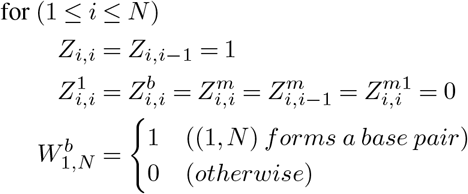

Recursion:
for (1 ≤ *i* < *j* ≤ *N*) s.t. (*j* − *i* ≤ *W*)

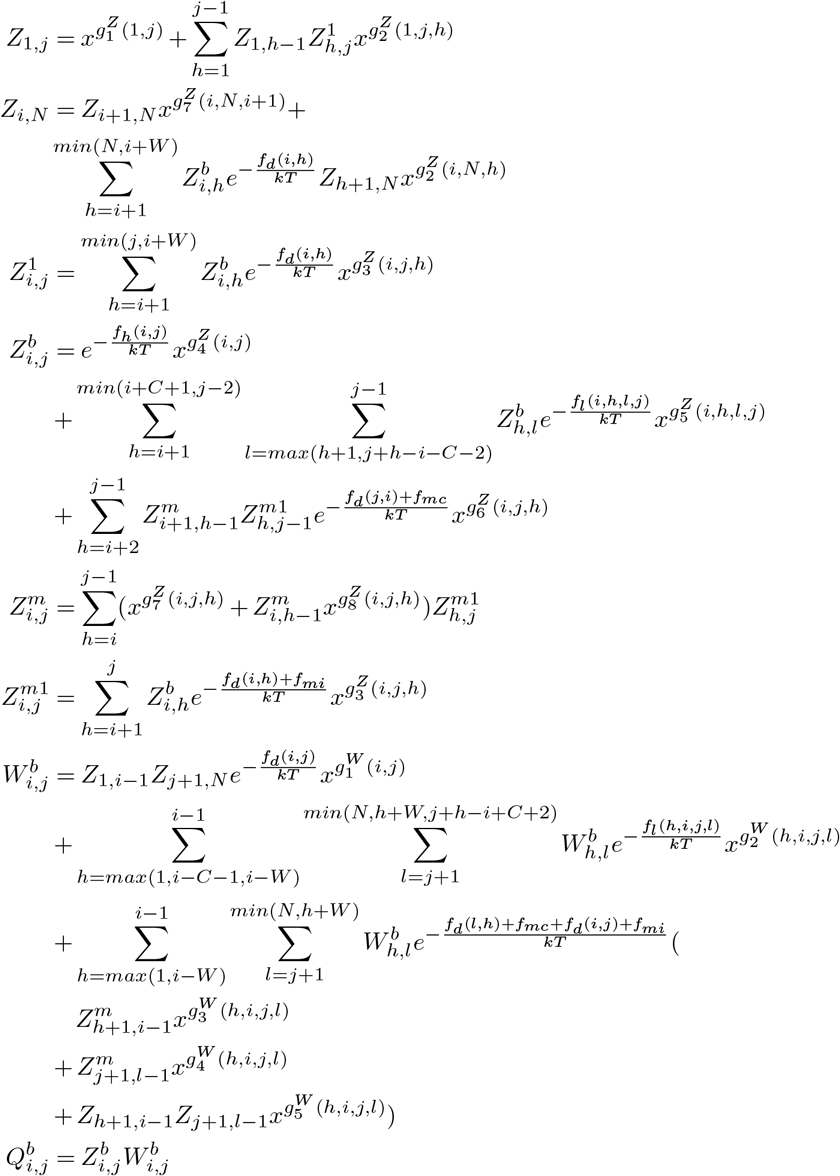

The *Z*_∗,∗_ functions are the *inside* partition functions, which represent the sums of all the Boltzmann factors in the corresponding sub-sequences. 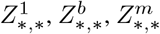, and 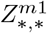 are the specified partition functions defined in the McCaskill algorithm (17). 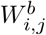 is the *outside* partition function, which represents the outside of the base-pair (*i*, *j*). The 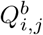 is the conditional partition function, the sum of all the Boltzmann factors when (*i*, *j*) forms a base pair.

The values 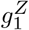 to 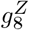 and 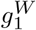 to 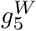, which are computed using the pre-computed function 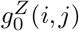, are the gains of the Hamming distance for the transitions represented by the recursions of partition functions. The significant difference from RintW is that the recursions of *Z*_1,*n*_, 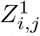, and 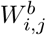 include the maximum-span constraint *W* of base-pairs in their range of the sum. A small improvement is that only edges are considered in the sum for *Z*_∗,∗_ (*Z*_1,*j*_ and *Z*_*i*,*N*_). Regarding the maximum-span constraint of base pairs, the algorithmic concept is equivalent to the calculation of dynamic programming (DP) variables *α*_*Outer*_ and *β*_*Outer*_ in Rfold (7) and ParasoR (11), but the notation of RintW is followed in the above recursions.

#### A.7. Fourier transform and post-processing

For all (*i*, *j*), such as (1 ≤ *i* ≤ *j* ≤ *N*) and (*j* − *i* ≤ *W*), by considering 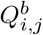 and *Z*_1,*N*_ of (*H*_*max*_ + 1) ways as a complex number sequence of (*H*_*max*_ + 1) elements and performing Fourier transformation, the conditional partition functions on each Hamming distance are obtained. Let *Z*(*d*)_1,*N*_ and 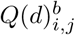 be the conditional partition functions on Hamming distance *d* of *Z*_1,*n*_ and 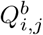, respectively. Then, the existence probability on Hamming distance *d* is written as

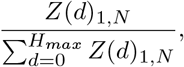

and the base-pairing probability on Hamming distance *d* is written as

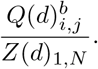

#### A.8. Computational complexity

In the following description, *N* is the length of the sequence, *H*_*max*_ is the maximum Hamming distance from the reference structure, and *U* is the degree of parallelism. Here *U* ≤ *H*_*max*_ + 1 is assumed. In the original RintW algorithm, the computational complexity of pre-processing is *O*(*N*^2^) in both time and space. In the partition function calculation, the time complexity is *O*(*N*^4^*H*_*max*_/*U*) and the space complexity is *O*(*N*^2^*H*_*max*_*U*). The original RintW uses DFT for post-processing; the time complexity of post-processing part is 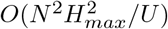, and its space complexity is *O*(*H*_*max*_*U*). Since *H*_*max*_ ≤ *N* holds, the computational complexity of the post-processing can be ignored in total complexity in both time and space. Finally, the time and space complexity of the original RintW algorithm as a whole are *O*(*N*^4^*H*_*max*_/*U*) and *O*(*N*^2^*H*_*max*_*U*), respectively.

When the maximum-span constraint is introduced, the computational complexity of pre-processing remains *O*(*N*^2^) in both time and space. In the distribution function calculation, the time complexity is *O*(*NW*^3^*H*_*max*_/*U*), and the space complexity is *O*(*NWH*_*max*_*U*) for the maximum-span of base pair *W*. When DFT is used for post-processing, the complexity of the post-processing is 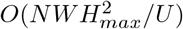 in time and *O*(*H*_*max*_*U*) in space. Because the *H*_*max*_ value may be close to *N*, the computational complexity of the post-processing cannot be ignored. By using FFT instead of DFT, we can reduce the time complexity of the post-processing component to *O*(*NW H*_*max*_*log*(*H*_*max*_)/*U*). Then, the total computational complexity is *O*(*N* (*N* + *WH*_*max*_(*W*^2^ + *log*(*H*_*max*_))/*U*)) in time and *O*(*N* (*N* + *WH*_*max*_*U*)) in space.

The summary of computational complexities is shown, with the notation simplified by using *H*_*max*_ ≤ *N*, in Table 1.

**Table 1.** Computational complexity of the existing and proposed methods are summarized. *N* = sequence length. *W* = maximum-span. Note that *H*_*max*_ ≤ *N* and *W* ≤ *N* always holds. *U* = degree of parallelism.

|  | RintW, time | RintC (proposed), time |
| --- | --- | --- |
| preprocessing | $O(N^2)$ | $O(N^2)$ |
| main calculation | $O(N^5/U)$ | $O(N^2W^3/U)$ |
| postprocessing (DFT) | $O(N^4/U)$ | $O(N^3W/U)$ |
| postprocessing (FFT) | $O(N^3 \log N/U)$ | $O(N^2W \log N/U)$ |
| total (DFT) | $O(N^5/U)$ | $O(N^2W(W^2 + N)/U)$ |
| total (FFT) | $O(N^5/U)$ | $O(N^2W(W^2 + \log N)/U)$ |
|  | RintW, space | RintC (proposed), space |
| preprocessing | $O(N^2)$ | $O(N^2)$ |
| main calculation | $O(N^3U)$ | $O(N^2WU)$ |
| postprocessing (DFT) | $O(NU)$ | $O(NU)$ |
| postprocessing (FFT) | $O(NU)$ | $O(NU)$ |
| total (DFT) | $O(N^3U)$ | $O(N^2WU)$ |
| total (FFT) | $O(N^3U)$ | $O(N^2WU)$ |

### B. Interval arithmetic and accuracy assurance

In this subsection we briefly explain the rounding mode control function of IEEE 754, and the accuracy assurance arithmetic. Representing real numbers by floating-point numbers can cause deviations from actual values. Therefore, numerical values can conceivably be held as an interval including the actual value. We define arithmetic operations between intervals to obtain an interval necessarily containing the results of arithmetic operations on actual values. Then, the upper bound of the numerical error is obtained as the width of the interval of the calculation result. Most modern computers use the IEEE 754 method for floating-point arithmetic. This method has a rounding mode control function, and we can specify truncation and rounding-up. By using this function, the accuracy assurance calculation described above can be executed efficiently.

Our accuracy assurance calculation used the kv library (18)) implemented in C++. The kv library is open source software and requires only C++ Boost for its backend.

### C. Logsumexp on complex numbers with interval arithmetic

A method to perform logsumexp on whole complex numbers has been developed. Details of the calculation algorithm are provided in the following subsections. There are different parts of algorithms for scalar and interval types, but those for scalar types are described in the supplementary file. In this subsection, only methods for interval types are described. If only the scalar type is considered, the complex number defined in polar coordinates and logsumexp defined only in terms of a radius are sufficient. Extensions to interval arithmetic, however, are complicated.

The Vienna RNA Package (19) prevents overflow by scaling. Their scaling factor construction is sophisticated, and under some assumptions, their scaling is equivalent to a kind of logsumexp. The original RintW (6) also utilized the same scaling technique as Vienna. However, with Vienna’s method, the deviation between the scaling factor and the value of the actual distribution function can increase exponentially, so overflowing cannot be completely avoided. Unlike them, logsumexp does not need scaling factors, and overflows are completely avoided.

#### C.1. Notation and representation

In this subsection, a bracketing character like [*x*] indicates an interval type variable. A pair of values in a bracket (e.g., [0, 1]) indicates a closed interval. When two variables are enclosed (e.g., [*x*, *y*]), each variable *x* and *y* is a scalar type (or floating-point type), not an interval type. It is possible to convert one scalar *x* into an interval type while guaranteeing accuracy. Such an interval variable is expressed as [*x*, *x*] (i.e., [*x*, *x*] is an interval that includes the real value *x*). Finally, a function *f*_*upper*_([*x*]) = *u* for obtaining the maximum value of the interval type variable [*x*] = [*l*, *u*], a function *f*_*lower*_([*x*]) = *l* for obtaining the minimum value, and a function 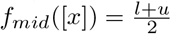 for obtaining the median value are used. It is assumed that they are not necessarily accuracy-guaranteed. In this subsection, the base of *log* is *e*.

To represent the complex number [*a*] + [*b*]*i*, ([*r*], [*c*], [*d*]) is held for

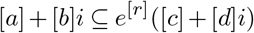

However, as a normalization condition,

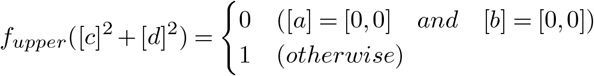

must be satisfied. It is assumed that 1 is numerically almost 1. The difference from 1 accumulates by multiplication, but it is reset by addition. For convenience, [*r*] = [0, 0] must be satisfied when ([*a*] = [0, 0] *and* [*b*] = [0, 0]).

The normalization protocol and conversion protocol between this and usual representation are described in the supplementary file. Multiplication and addition protocols are described below.

#### C.2. Multiplication

The multiplication of the two values ([*r*_1_], [*c*_1_], [*d*_1_]) and ([*r*_2_], [*c*_2_], [*d*_2_]) can be described as

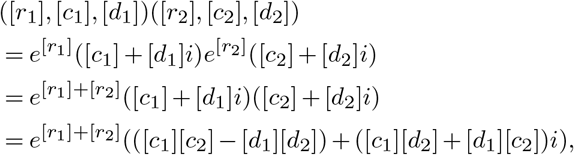

and ([*r*_1_] + [*r*_2_], [*c*_1_][*c*_2_] − [*d*_1_][*d*_2_], [*c*_1_][*d*_2_] + [*d*_1_][*c*_2_]) is obtained as a solution. In normalization post-processing, if [*c*_1_][*c*_2_] − [*d*_1_][*d*_2_] = [*c*_1_][*d*_2_] + [*d*_1_][*c*_2_] = [0, 0], [*r*] = [0, 0] is substituted. Otherwise, because the product of the complex numbers with absolute value 1 is absolute value 1, it is naturally normalized.

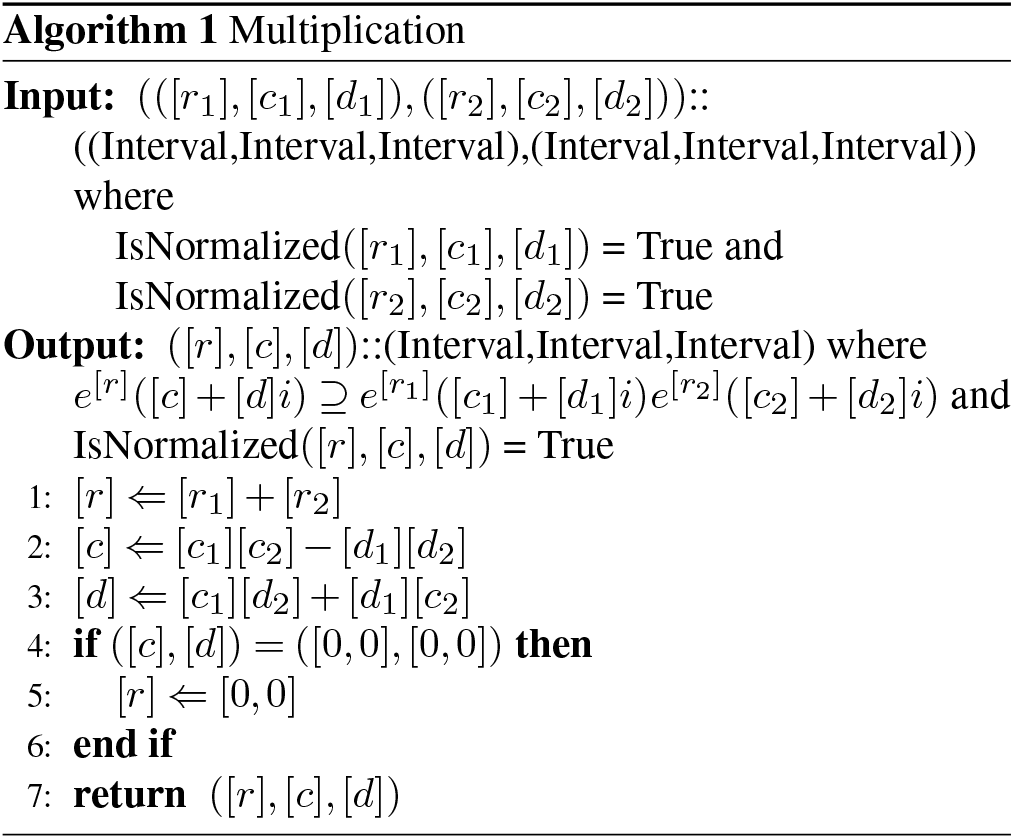

#### C.3. Addition

Consider the sum of the two values ([*r*_1_], [*c*_1_], [*d*_1_]) and ([*r*_2_], [*c*_2_], [*d*_2_]). Since addition is commutative, assuming *f_mid_*([*r*_1_]) ≥ *f*_*mid*_([*r*_2_]) does not decrease generality. Then, it can be formulated as

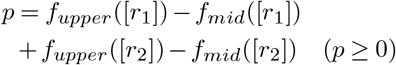

and

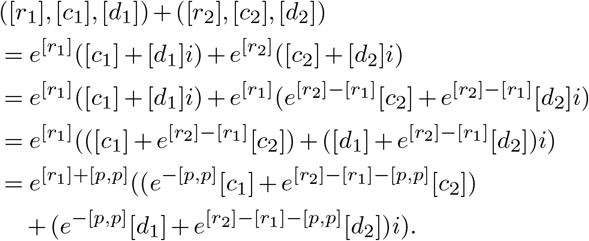

Thus, *f*_*upper*_(*e*^−[*p*,*p*]^) ≤ 1 follows from the assumption of *p* ≥ 0. Additionally, 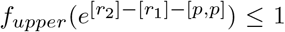 follows from the assumption that *f*_*mid*_([*r*_1_]) ≥ *f*_*mid*_([*r*_2_]) (the proof is provided in the Supplementary materials). Therefore *e*^−[*p*,*p*]^ and 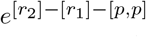 can be directly calculated without overflow occurring. Therefore,

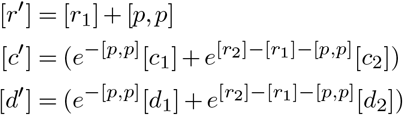

can be calculated, and ([*r*′], [*c*′], [*d*′]) satisfies

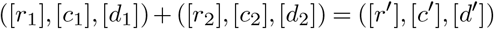

as the summation. Finally, since this is not normalized, normalization processing is required.

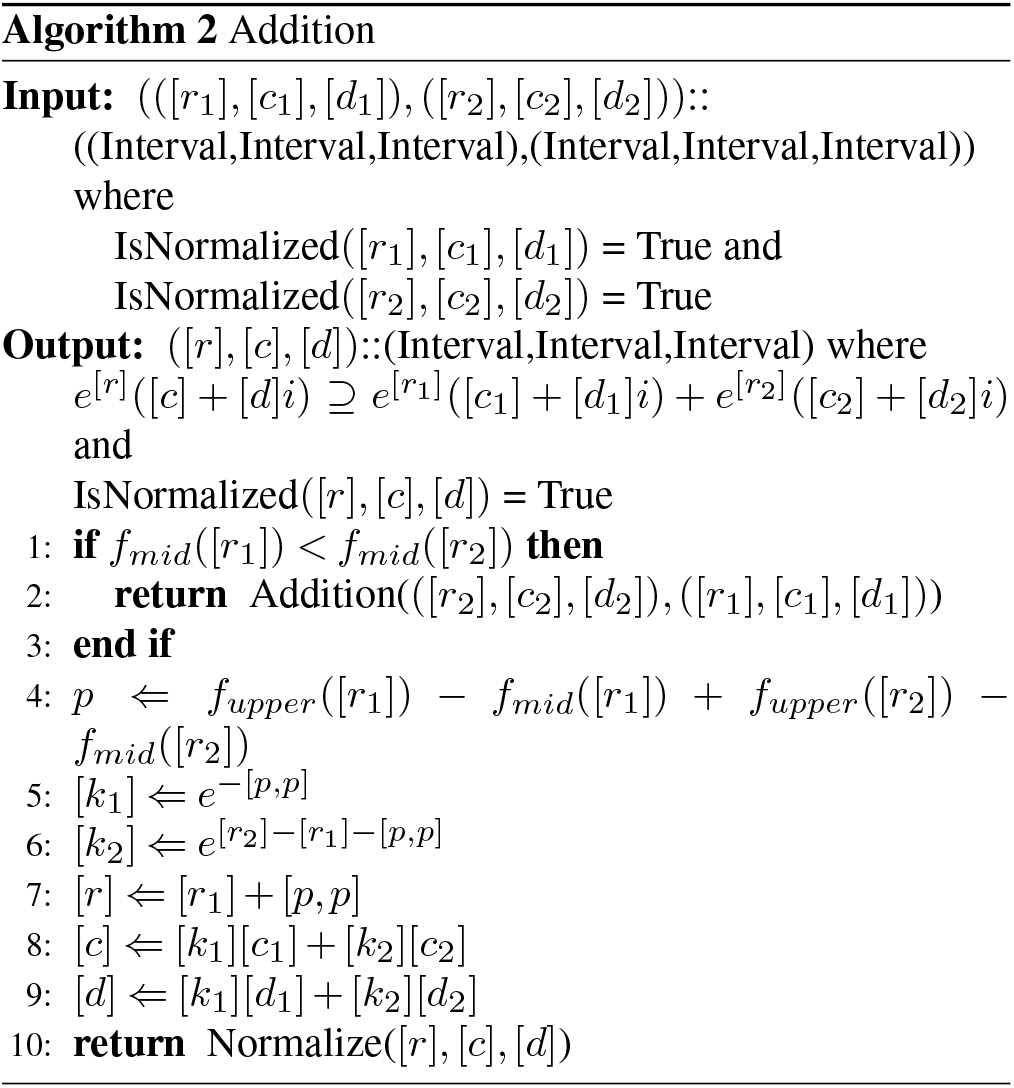

In the classic logsumexp, it is known that one can reduce the numerical error of summation by using summation-specific technique than recursively using the two-operand addition function. For the summation-specific technique, in three or more operands, one choice the maximum number as the scaling factor and scale the others. On the other hand, we developed only the normal two-operand addition function. Following experiment showed that our method brings sufficient numerical accuracy. Nevertheless, further improvement may be still possible.

#### C.4. Requirements for an accuracy assurance calculation library

The functions that the accuracy assurance calculation library must perform in this method are as follows:

1. The conversion from scalar to interval type guarantees accuracy.
2. Four arithmetic operations, log, and exp, with accuracy assurance for the interval type.
3. The previously described *f*_*upper*_([*x*]) and *f*_*mid*_([*x*]).

## Results and Discussion

### D. Experimental procedure

The S151 Rfam Dataset ‘with all pseudoknots removed’ (20) was used for evaluation of time complexity and numerical accuracy in interval operation.

For the application of our proposed method to RNA molecules longer than those in the S151 Rfam dataset (20), the primary sequences and the corresponding native secondary structures of 16S rRNAs were obtained from three-dimensional structures of *E. coli* and *T. thermophilus* 70S ribosomes from the Nucleic Acid Database (NDB, (21), (22). The NDB IDs of the *E. coli* and *T. thermophilus* ribosome structures were 4V9D (chainID: AA)(23) and 4V51 (chainID: AA)(24), respectively. As the secondary structures of these 16S rRNAs, base pairs were selected according to the "base pair hydrogen bonding classification" provided by NDB. Specifically, base-pairs were classified as 1 in the Leontis–Westhof classification (25) and either 19, 20, or 28 in the Saenger classification (26). Base-to-base correspondence between the primary sequence and its secondary structure (derived from the three-dimensional structure in which several residues are missing) was estimated using Needleman-Wunsch alignment (27).

The energy parameter rna_turner2004.par included in the Vienna RNA package (19) version 2.4.9 was used. However, the source code itself of Vienna was not used. The algorithms were implemented by the authors, except for parameter file reading, which is based on ParasoR’s implementation. (11)

### E. Computation time

To demonstrate computational efficiency, the computation time of the proposed method using the S151 Rfam Dataset (20) was measured. In the proposed method, the reference structures were obtained by Centroid-Fold (28)(*γ* = 1.0). All cores of the Intel Core i7 4770 CPU were used as a computational resource in this measurement.

We measured the computation time in the case where the maximum-span constraint *W* = 100 is introduced and in the case where no restriction is applied (equivalent to RintW (6) (Figure 1). This result shows the computation time of the methods using the constraint *W* = 100 and no-restriction roughly, which follow the theoretical complexities of the square and fifth power, respectively, confirming that computational complexity can be reduced by introducing the maximum-span constraint into the proposed method.

**Fig. 1.**
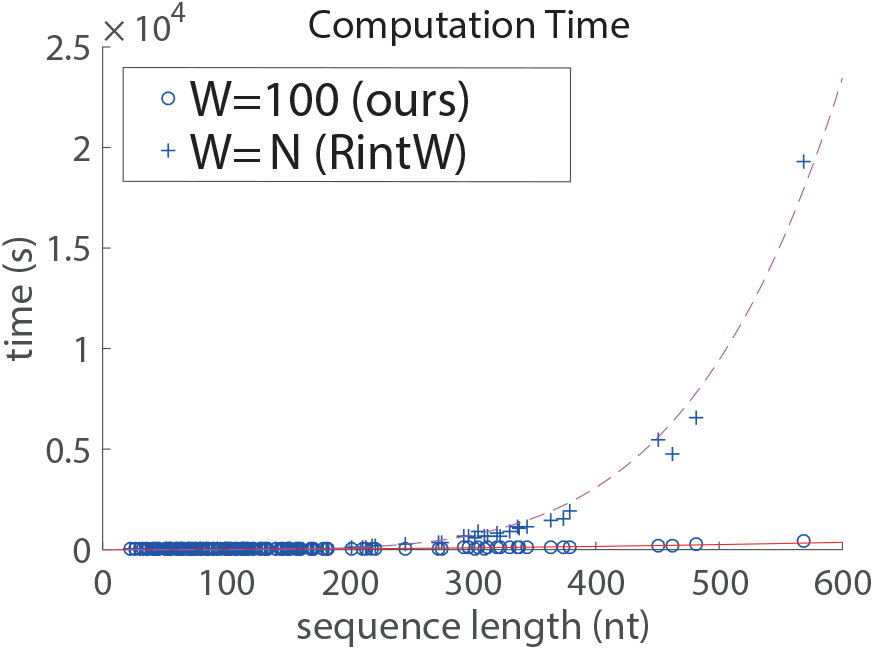
Calculation time of the proposed method. The red solid line represents *y* = 0.0010101*x*^2^. The purple dashed line represents *y* = 3.0163*e* − 10*x*^5^. Both lines were obtained by fitting to the result using the least squares method.

We also examined how the calculation time changes when the value of the maximum-span constraint *W* is changed (Figure 2). 32 Cores of the Intel Xeon Gold 6130 were used as a computational resource in this experiment. As a result, when *N* ≤ *W*, the same calculation time is required regardless of the value of *W*. When *W* < *N*, the calculation time is reduced by the effect of the maximum-span.

**Fig. 2.**
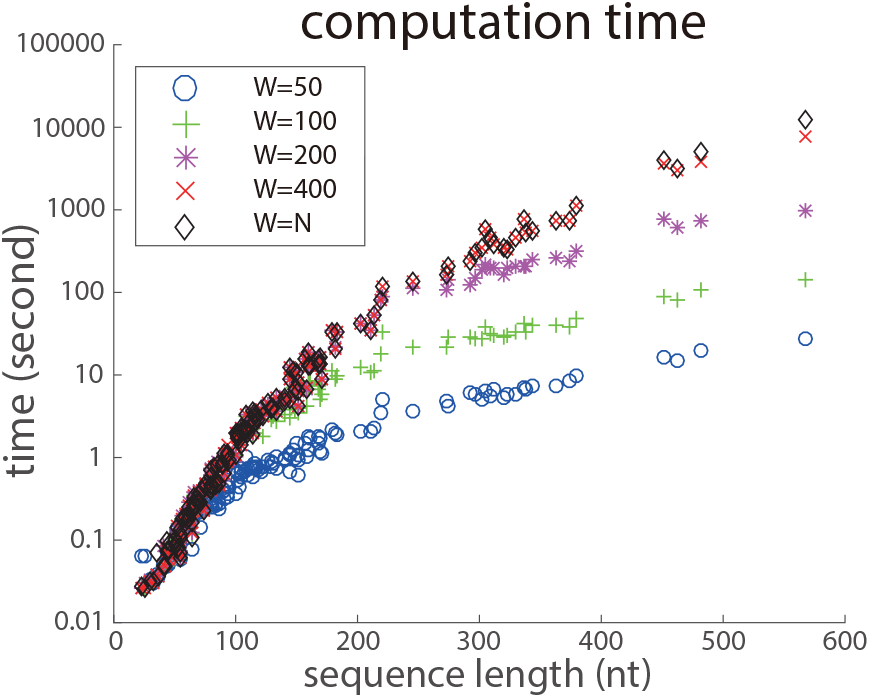
Calculation time of the proposed method. Each data point is the calculation time of a single sequence from S151Rfam dataset.

### F. Thermal stability of ribosomal RNA

As an application of the proposed method, we analyzed the thermal stability of the secondary structures of 16S rRNAs derived from *E. coli* and *T. thermophilus* using Credibility Limit (29) as a metric. Credibility Limit is a distance in which a certain percentage of structures is distributed. The larger the Credibility Limit value, the more intense the thermal fluctuation of the molecule. The Credibility Limit (0.5) was obtained from the Hamming distances of the secondary structures under the proposed method with temperatures ranging from 37 to 55 degrees Celsius (Figure 3). For the 16S rRNAs, three types of reference structure were prepared. (i) The “initial reference” was obtained using CentroidFold. (ii) The “refined reference” was obtained by the following steps: RintC was performed with the initial reference; a region whose existence probability is locally high was chosen; BPPM of the region was calculated; and CentroidFold was used to determine a “refined reference” from the BPPM. (iii) The “natural reference” was the reference structure derived from the three-dimensional structure in NDB. This result shows that the rRNA of thermophilic bacteria *T. thermophilus* had a lower Credibility Limit than that of *E. coli*, which suggests that the thermal stability of *T. thermophilus* rRNA is higher than that of *E. coli* rRNA. According to an *in vivo* assay (30), thermophilic bacteria have reduced dynamics of intracellular macromolecules compared with mesophilic bacteria. Our experimental result is consistent with their results. The use of the natural secondary structure as a representative structure exhibited a relatively higher Credibility Limit, compared with the “initial” and “refined” references. This implies the DP calculation with the Turner model is compatible for the representative structure derived from the Turner model, such as the “initial” and “refined” references. Note that this result would not indicate the advantage of “initial” and “refined” references over the “natural” reference.

**Fig. 3.**
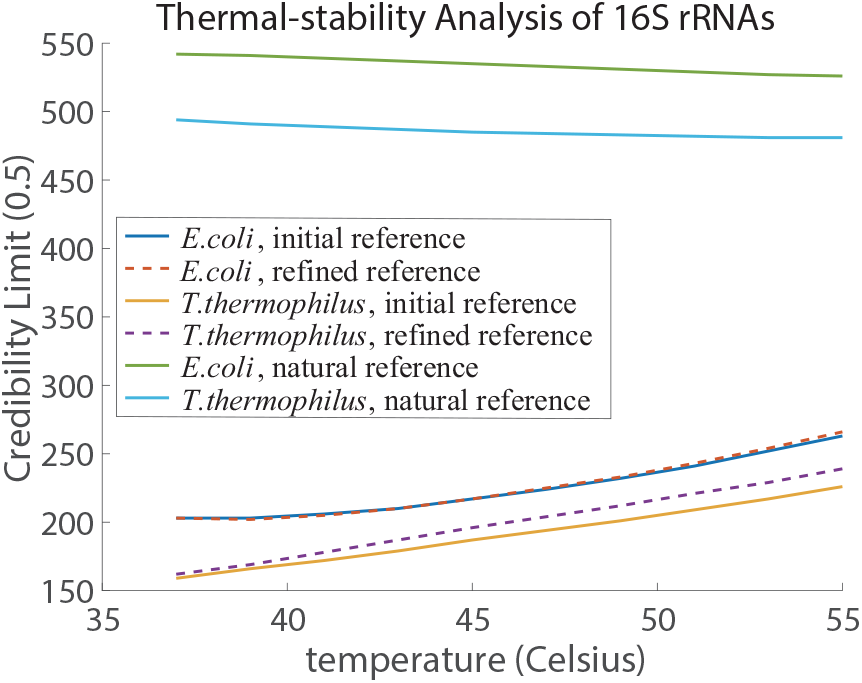
Thermal-stability analysis for secondary structures of *E. coli* and *T. thermophilus* 16S rRNAs. The “initial reference” is the reference structure obtained using CentroidFold. The “refined reference” is the reference structure obtained using RintC and the base-pairing probability matrix (BPPM) (see the section “Thermal stability of ribosomal RNA” for the details). The “natural reference” is the reference structure derived from the three-dimensional structure. The “experimental procedure” section provides a detailed description about the “natural reference.”

### G. Numerical error evaluation

For a quantitative evaluation of RintC numerical error, accuracy-guaranteed numerical computations with interval arithmetic were applied to the calculation process of RintC with the RF00008B sequence in the S151 Rfam dataset (20). The length of the RF00008B sequence was short enough for the evaluation of time-consuming calculation without any type of Fourier transform. The numerical errors of three types of calculation (DFT, FFT, and non-FFT) are shown in Figure 4. For the DFT and FFT methods, in this result, the numerical error (i.e., interval width) is almost equal to 1, when the calculated existence probability is quite small. Interval width = 1 indicates that the probability is within [0, 1], provides no meaningful information owing to numerical error. In contrast, the numerical error remains low when the existence probability is moderate or high. DFT-based results are slightly more accurate than FFT-based results. In further numerical error comparisons between non-Fourier transform results and DFT or FFT results, the numerical error of the non-Fourier result is smaller than those of the DFT or FFT results. This implies that the problematic numerical error is indeed caused by Fourier transform.

**Fig. 4.**
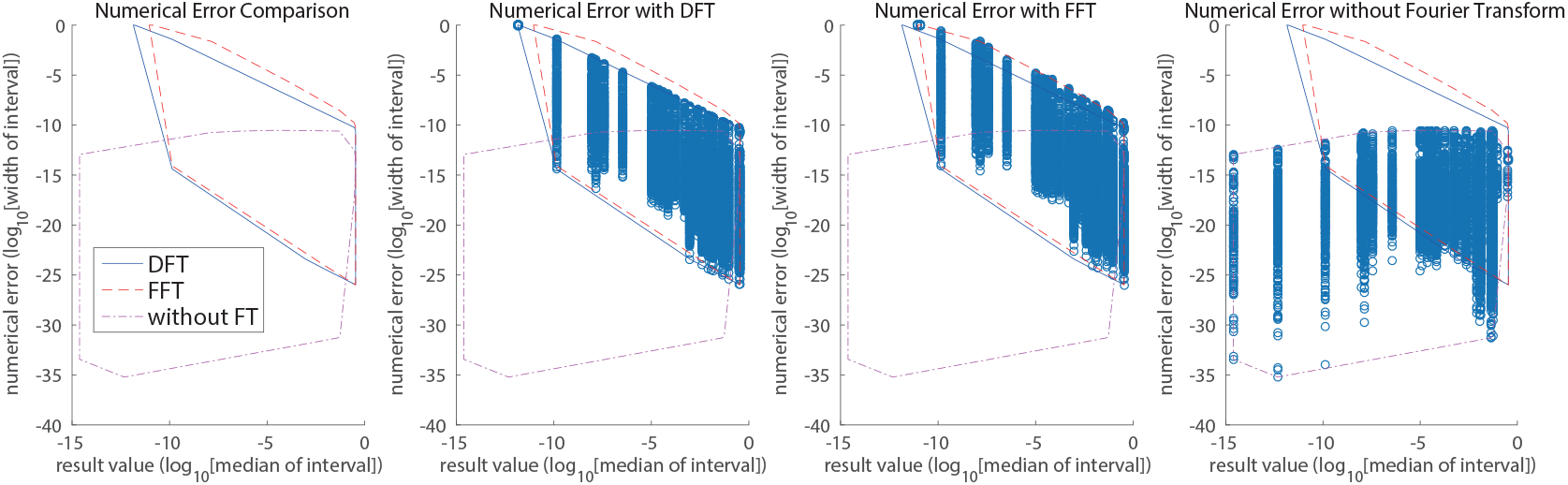
The result of the numerical error experiment with RF00008B(54 nucleotides). The leftmost plot explains the convex hulls for the result values and their errors under each experimental condition. The three plots to the right are scatter plots of the raw data for the result values and errors under one experimental condition and the convex hulls for each. In this evaluation, the reference structure was obtained by CentroidFold.

The numerical error evaluation was also conducted for all sequences in the S151 Rfam dataset and *E. coli* 16s rRNA (Figure 5). Each data point corresponds to an individual sequence in the S151 Rfam dataset or *E. coli* 16s rRNA. In comparisons of the numerical errors between DFT and FFT versions (Figure 5a), DFT is always more accurate than FFT. This result is consistent with that shown in Figure 4. In addition, a relationship between the numerical error and sequence length in the DFT results was also investigated (Figure 5b). This result demonstrates that the numerical error of 16S rRNA is almost equal to −7, which suggests that the numerical error for long RNA sequences (≥ 1000 nt) is sufficiently small for structures with a moderate or high probability of existence. This accuracy is sufficient for thermal stability analysis because an accurate evaluation of large clusters is only required for their analysis.

**Fig. 5.**
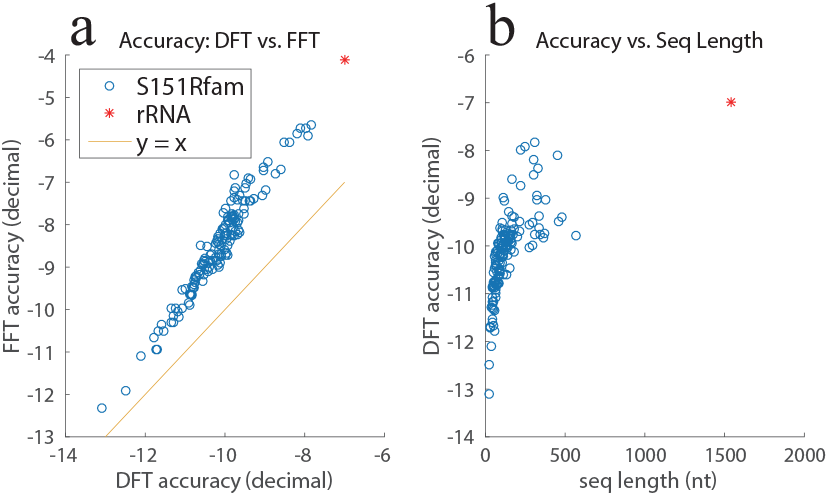
The results of the numerical error experiment. (a) Numerical error comparison. (b) Relationship between sequence length and numerical error. The *x* and *y* axes have minima of (*log*_10_ (*median*) + *log*_10_ (*width*)) under the DFT and FFT methods. The reference structure was obtained using CentroidFold.(28) (γ =1.0)

## Conclusions

Since RNA secondary structures have large thermal fluctuations, prediction of the most stable secondary structure is insufficient for representing native structural behavior of an RNA molecule. Marginal probabilities on Hamming distances from reference structures, which represent the landscape of all the possible RNA secondary structures, can be efficiently computed by combining Fourier transform with dynamic programming, but the computational costs are still too high for long RNAs.

In this research, we have implemented a maximum-span constraint of base pairs to reduce computational complexity. For long RNAs, however, there remains another problem: numerical overflow. Since the standard method for avoiding overflow in stochastic models, logsumexp (logarithm of sum of exponentials), is not directly applicable to Fourier transform, we have developed an extended logsumexp method for whole complex numbers. We have shown that reduced computational time enables us to analyze the thermal-stability of long RNAs, such as 16S ribosomal RNAs.

We have also adopted accuracy-guaranteed numerical computation with interval arithmetic to evaluate numerical errors. We have shown that numerical errors for small probabilities are substantial when FFT or DFT is used. Quantitative assessment of the observed numerical instabilities, however, reveal that our method achieves sufficient numerical accuracy for thermodynamic stability analysis of RNA secondary structures. These results demonstrate that our method is a powerful tool for understanding long RNAs.

## ACKNOWLEDGEMENTS

The authors deeply thank the developers of the RintW, ParasoR, and ViennaRNA Package implementations, which were necessary for the developed software tool. The authors also deeply thank Dr. Shun Sakuraba, who suggested the possible application of maximum-span constraint and numerical problems in previous research. The authors also thank members of the Artificial Intelligence Research Center, AIST, and Hisanori Kiryu’s laboratory for useful discussions. Computations were partially performed on the NIG supercomputer at the ROIS National Institute of Genetics.

This work was supported by MEXT/JSPS KAKENHI Grant Numbers JP16H02484 and JP16H06279 to K.A. and JP16K16143 to J.I, JST CREST Grant Number JP-MJCR18S1 to K.A.

## Supplementary Note 1: All Functions for Logsumexp on Complex Number but not Interval

Here we explain how to perform logsumexp on the whole complex number. In this subsection, we describe methods for the normal scalar type (or floating point number type) which is not an interval type. In this section, the base of *log* is *e*.

### A. representation

First, as a representation of the complex number *a* + *bi*, hold (*r*, *c*, *d*) of

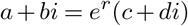

However, as a normalization condition,

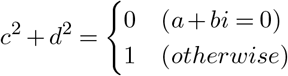

must be satisfied. For convenience, it must be satisfied that *r* = 0 when *a* + *bi* = 0. The algorithm for checking whether (*r*, *c*, *d*) is normalized can be written as follows.

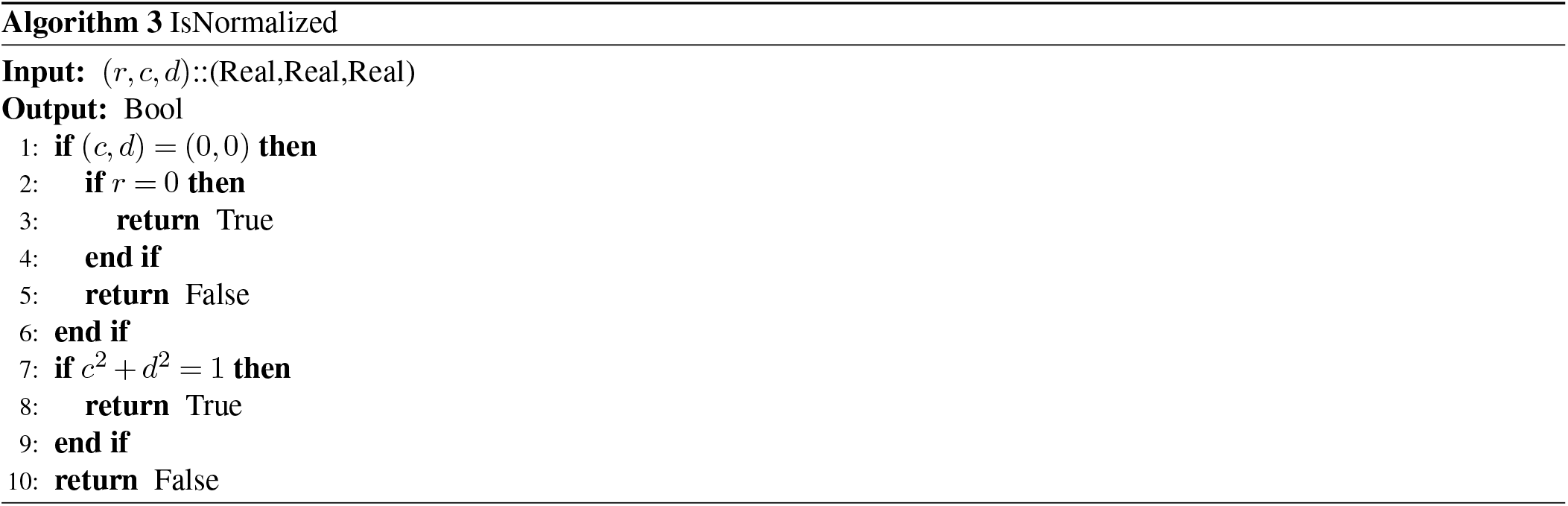

### B. normalization

When a number (*r*′, *c*′, *d*′) which is not normalized is given, a method of obtaining the normalized number (*r*, *c*, *d*) = (*r*′, *c*′, *d*′) is as follows.

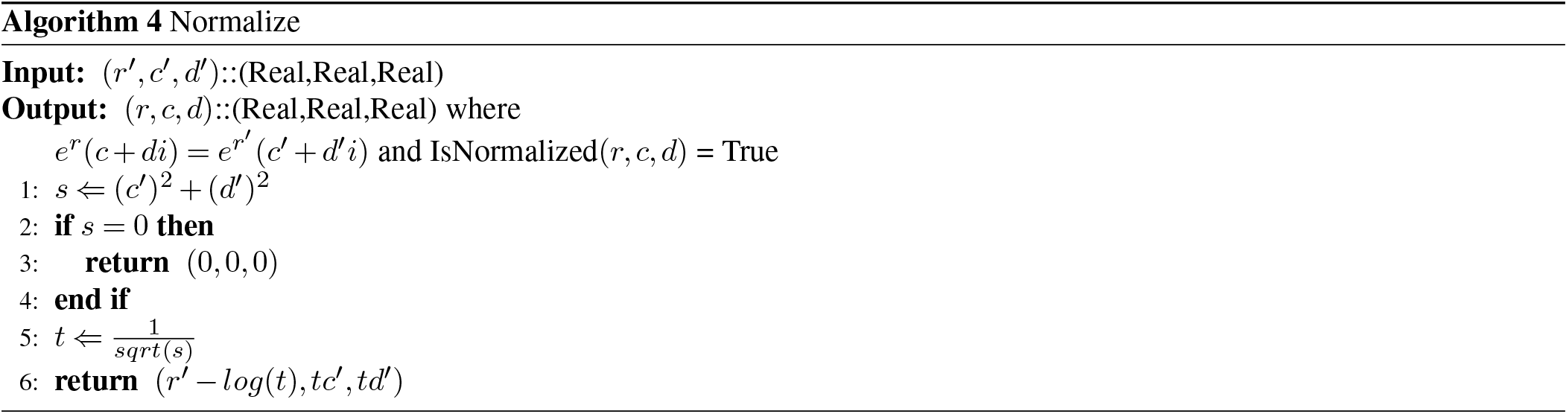

Description:

First, compute

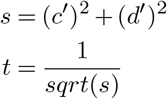

*t* is the reciprocal of the absolute value of the input. At this time

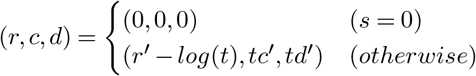

is a normalized solution.

### C. conversion

To obtain (*r*, *c*, *d*) when (*a*, *b*) such as *a* + *bi* is given, normalize (0, *a*, *b*).

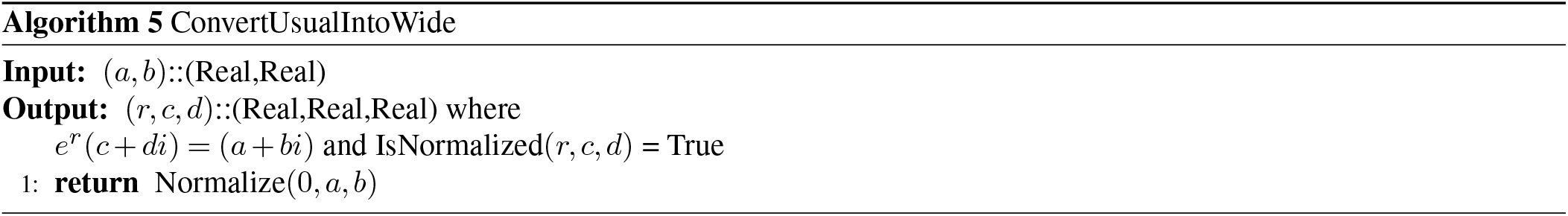

For inverse transformation, calculate normally.

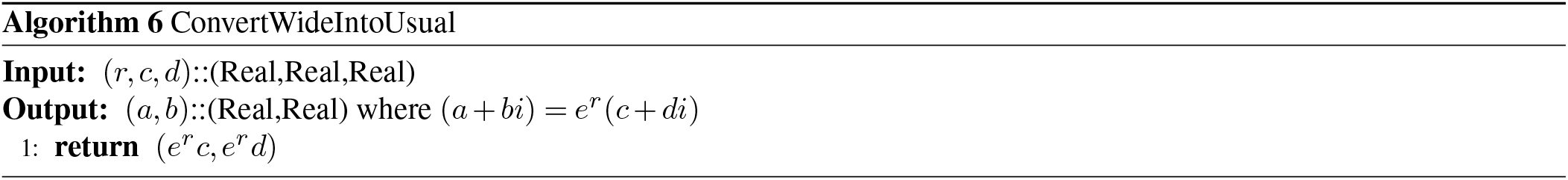

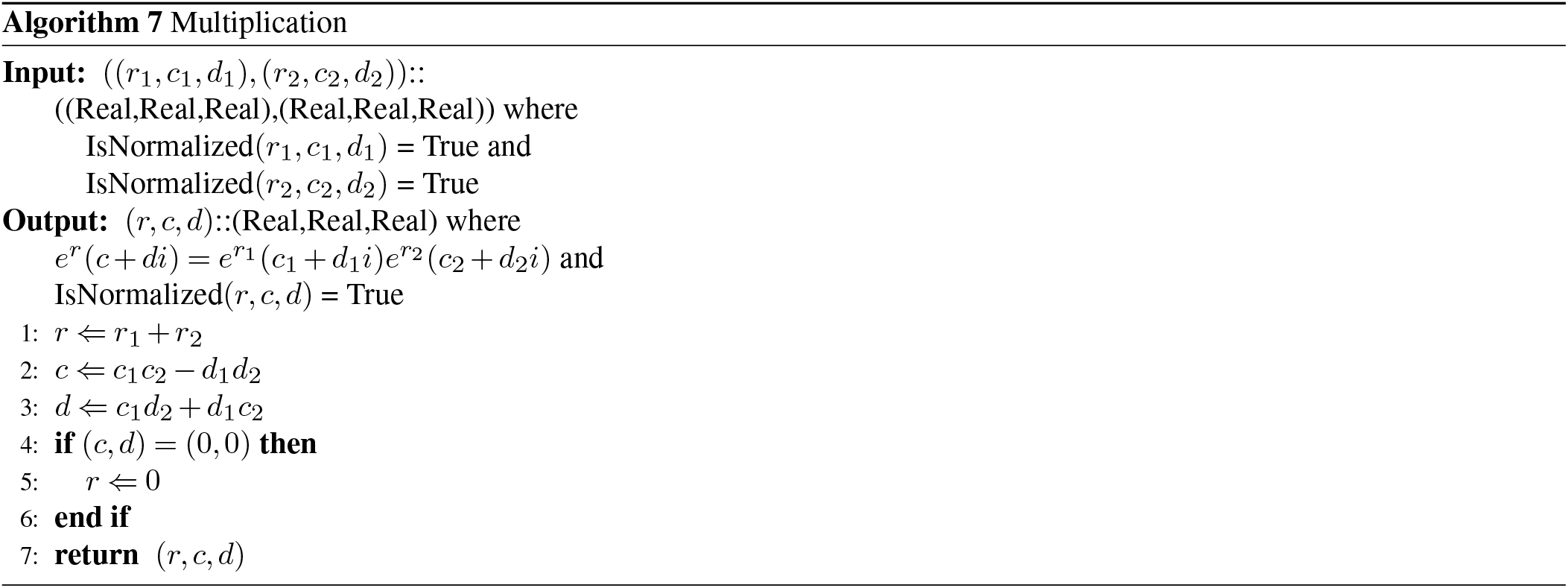

### D. multiplication

Description:

The multiplication of the two values (*r*_1_, *c*_1_, *d*_1_) and (*r*_2_, *c*_2_, *d*_2_) can be descripted as

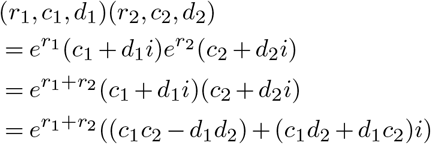

and (*r*_1_ + *r*_2_, *c*_1_*c*_2_ − *d*_1_*d*_2_, *c*_1_*d*_2_ + *d*_1_*c*_2_) is obtained as a solution. As a post-processing for normalization, if *c*_1_*c*_2_ − *d*_1_*d*_2_ = *c*_1_*d*_2_ + *d*_1_*c*_2_ = 0, substitute *r* = 0. Otherwise, since the product of the complex numbers with absolute value 1 is absolute value 1, it is naturally normalized.

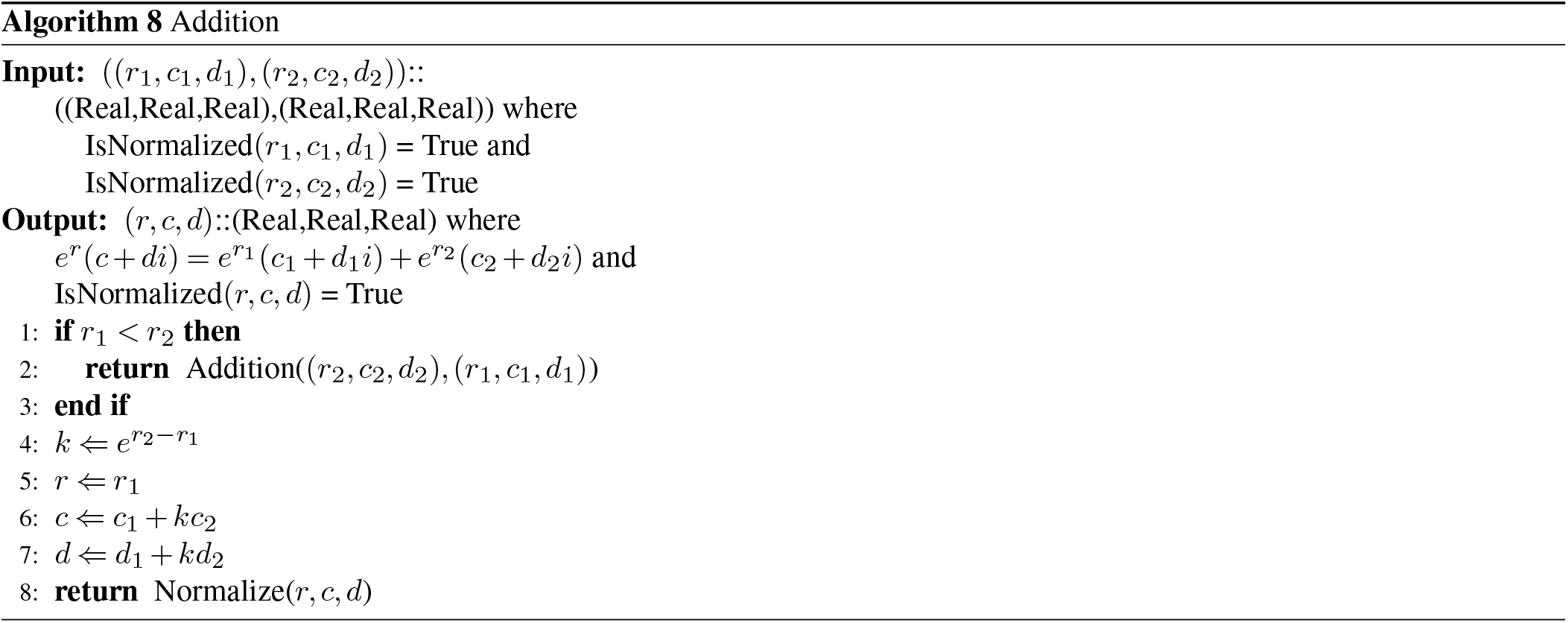

### E. addition

Description:
Consider adding the two values (*r*_1_, *c*_1_, *d*_1_) and (*r*_2_, *c*_2_, *d*_2_). Since addition is commutative, assuming *r*_1_ ≥ *r*_2_ does not lose generality. Then, it can be formulated as

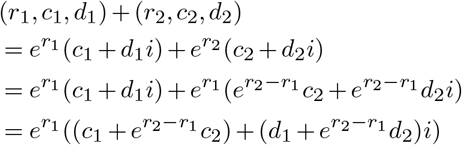

Since it is 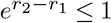 from the assumption of 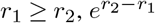 can be directly calculated without overflowing. Therefore,

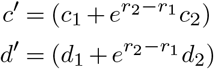

can be calculated and (*r*_1_, *c*′, *d*′) satisfies

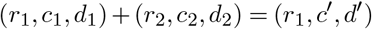

as an answer of addition. Finally, since this is not normalized, it needs normalization processing.

## Supplementary Note 2: Auxilary Functions for Logsumexp on Complex Number and Interval

The algorithm for checking whether ([*r*], [*c*], [*d*]) is normalized can be written as follows.

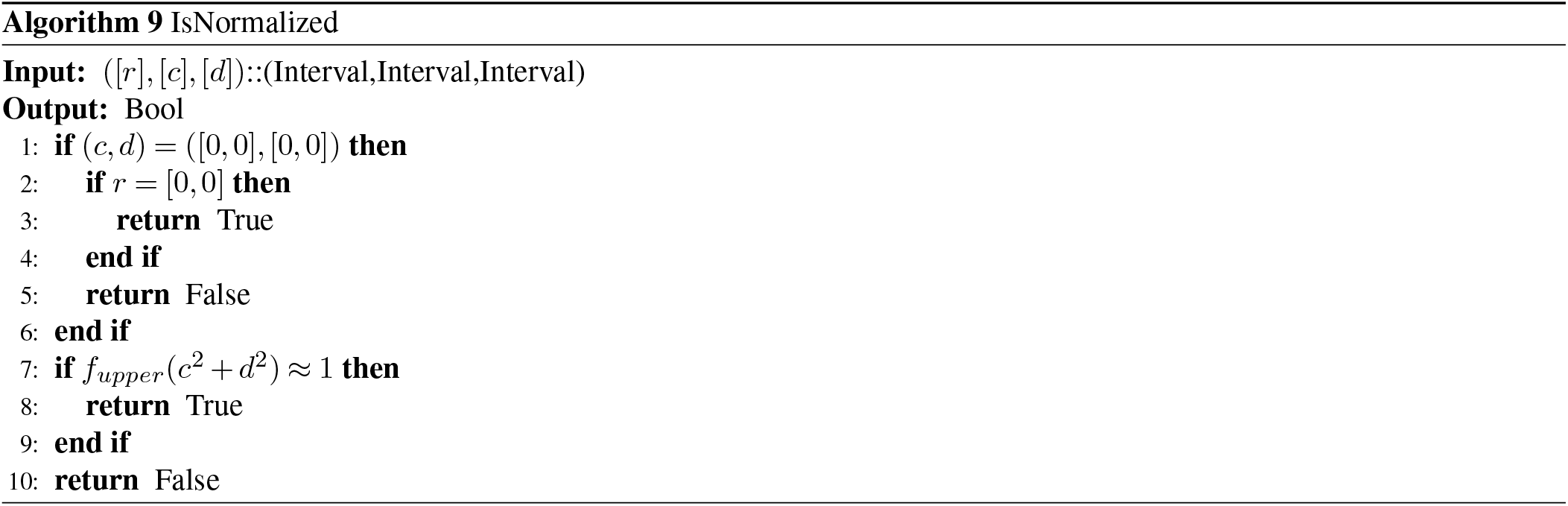

### A. normalization

When a number ([*r*′], [*c*′], [*d*′]) which is not normalized is given, a method of obtaining the normalized number with accuracy assurance ([*r*], [*c*], [*d*]) ⊇ ([*r*′], [*c*′], [*d*′]) is as follows.

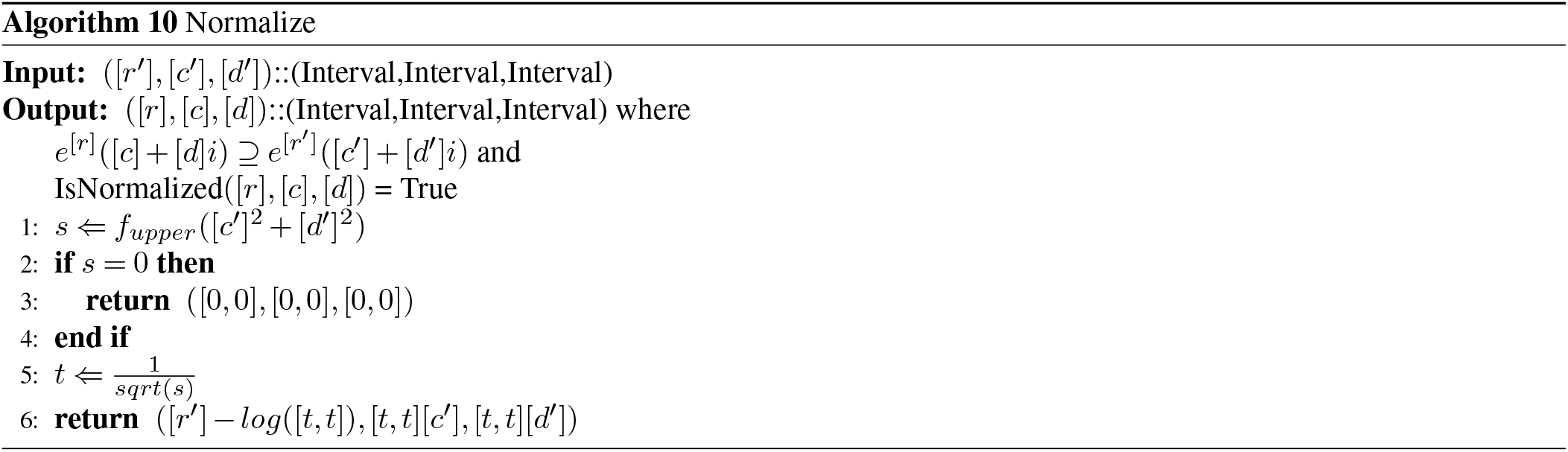

Description:

First, compute

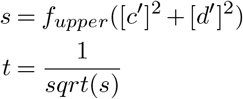

*t* is the reciprocal of maximum value of the absolute value of the input. At this time

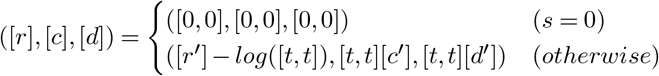

is a normalized solution.

### B. conversion

To obtain ([*r*], [*c*], [*d*]) when [*a*] + [*b*]*i* is given, normalize ([0, 0], [*a*], [*b*]).For inverse transformation, calculate normally.

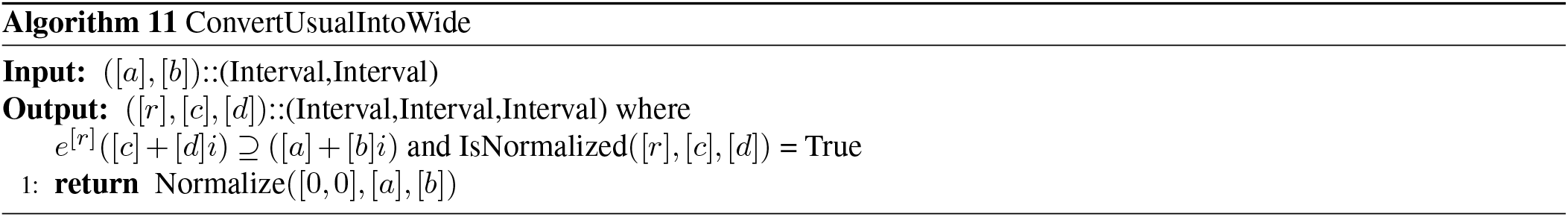

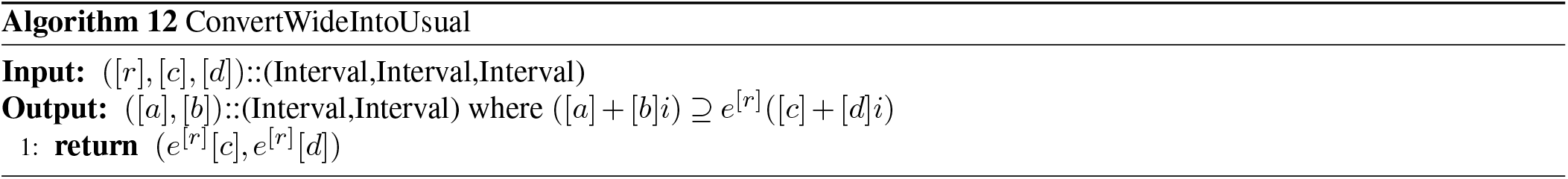

## Supplementary Note 3: Proof of Proposition 1

Proposition 1: 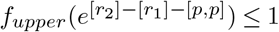

Proof:

Since they are assumptions with *f*_*mid*_([*r*_1_]) ≥ *f*_*mid*_([*r*_2_]) and

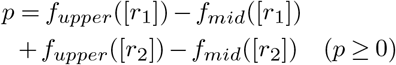

it holds that

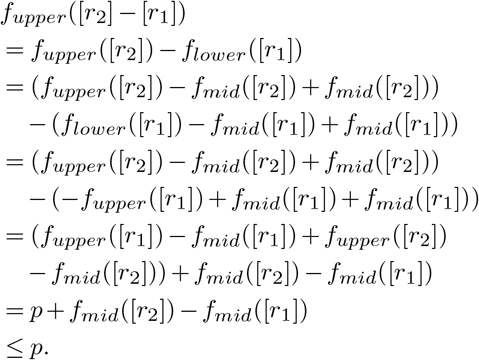

Therefore,

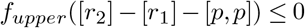

holds. Since,

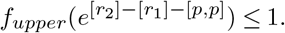

QED.

## Bibliography

1. E. Freyhult, V. Moulton, and P. Clote. RNAbor: a web server for RNA structural neighbors. Nucleic Acids Research, 35(Web Server):W305–W309, may 2007. ISSN 0305-1048. doi: 10.1093/nar/gkm255.

2. Ronny Lorenz, Christoph Flamm, and Ivo L. Hofacker. 2D projections of RNA folding landscapes. IN: GERMAN CONFERENCE ON BIOINFORMATICS, pages 11–20, 2009.

3. Lee A. Newberg and Charles E. Lawrence. Exact Calculation of Distributions on Integers, with Application to Sequence Alignment. Journal of Computational Biology, 16(1):1–18, jan 2009. ISSN 1066-5277. doi: 10.1089/cmb.2008.0137.

4. Evan Senter, Saad Sheikh, Ivan Dotu, Yann Ponty, and Peter Clote. Using the Fast Fourier Transform to Accelerate the Computational Search for RNA Conformational Switches. PLoS ONE, 7(12):e50506, dec 2012. ISSN 1932-6203. doi: 10.1371/journal.pone.0050506.

5. Ryota Mori, Michiaki Hamada, and Kiyoshi Asai. Efficient calculation of exact probability distributions of integer features on RNA secondary structures. BMC Genomics, 15(Suppl 10):S6, dec 2014. ISSN 1471-2164. doi: 10.1186/1471-2164-15-S10-S6.

6. Taichi Hagio, Shun Sakuraba, Junichi Iwakiri, Ryota Mori, and Kiyoshi Asai. Capturing alternative secondary structures of RNA by decomposition of base-pairing probabilities. BMC Bioinformatics, 19(S1):38, feb 2018. ISSN 1471-2105. doi: 10.1186/s12859-018-2018-4.

7. Hisanori Kiryu, Taishin Kin, and Kiyoshi Asai. Rfold: an exact algorithm for computing local base pairing probabilities. Bioinformatics, 24(3):367–373, feb 2008. ISSN 1460-2059. doi: 10.1093/bioinformatics/btm591.

8. Evan Senter, Ivan Dotu, and Peter Clote. RNA folding pathways and kinetics using 2D energy landscapes. Journal of Mathematical Biology, 70(1-2):173–196, jan 2015. ISSN 0303-6812. doi: 10.1007/s00285-014-0760-4.

9. Juraj Michálik, Hélène Touzet, and Yann Ponty. Efficient approximations of RNA kinetics landscape using non-redundant sampling. Bioinformatics, 33(14):i283–i292, jul 2017. ISSN 1367-4803. doi: 10.1093/bioinformatics/btx269.

10. Teruo Sunaga. Theory of an interval algebra and its application to numerical analysis. Japan Journal of Industrial and Applied Mathematics, 26(2-3):125–143, oct 1958. ISSN 0916-7005. doi: 10.1007/BF03186528.

11. Risa Kawaguchi and Hisanori Kiryu. Parallel computation of genome-scale RNA secondary structure to detect structural constraints on human genome. BMC Bioinformatics, 17(1): 203, dec 2016. ISSN 1471-2105. doi: 10.1186/s12859-016-1067-9.

12. Hillary S.W. Han and Christian M. Reidys. The 5-3 Distance of RNA Secondary Structures. Journal of Computational Biology, 19(7):867–878, jul 2012. ISSN 1066-5277. doi: 10.1089/cmb.2011.0301.

13. David H Mathews, Matthew D Disney, Jessica L Childs, Susan J Schroeder, Michael Zuker, and Douglas H Turner. Incorporating chemical modification constraints into a dynamic programming algorithm for prediction of RNA secondary structure. Proceedings of the National Academy of Sciences of the United States of America, 101(19):7287–92, may 2004. ISSN 0027-8424. doi: 10.1073/pnas.0401799101.

14. I. L. Hofacker, B. Priwitzer, and P. F. Stadler. Prediction of locally stable RNA secondary structures for genome-wide surveys. Bioinformatics, 20(2):186–190, jan 2004. ISSN 1367-4803. doi: 10.1093/bioinformatics/btg388.

15. S. H. Bernhart, I. L. Hofacker, and P. F. Stadler. Local RNA base pairing probabilities in large sequences. Bioinformatics, 22(5):614–615, mar 2006. ISSN 1367-4803. doi: 10.1093/bioinformatics/btk014.

16. Sita J. Lange, Daniel Maticzka, Mathias Möhl, Joshua N. Gagnon, Chris M. Brown, and Rolf Backofen. Global or local? Predicting secondary structure and accessibility in mRNAs. Nucleic Acids Research, 40(12):5215–5226, jul 2012. ISSN 1362-4962. doi: 10.1093/nar/gks181.

17. J. S. McCaskill. The equilibrium partition function and base pair binding probabilities for RNA secondary structure. Biopolymers, 29(6-7):1105–1119, may 1990. ISSN 0006-3525. doi: 10.1002/bip.360290621.

18. Masahide Kashiwagi. kv - a C++ Library for Verified Numerical Computation, 2018.

19. Ronny Lorenz, Stephan H Bernhart, Christian Höner zu Siederdissen, Hakim Tafer, Christoph Flamm, Peter F Stadler, and Ivo L Hofacker. ViennaRNA Package 2.0. Algorithms for Molecular Biology, 6(1):26, nov 2011. ISSN 1748-7188. doi: 10.1186/1748-7188-6-26.

20. C. B. Do, D. A. Woods, and S. Batzoglou. CONTRAfold: RNA secondary structure prediction without physics-based models. Bioinformatics, 22(14):e90–e98, jul 2006. ISSN 1367-4803. doi: 10.1093/bioinformatics/btl246.

21. Buvaneswari Coimbatore Narayanan, John Westbrook, Saheli Ghosh, Anton I. Petrov, Blake Sweeney, Craig L. Zirbel, Neocles B. Leontis, and Helen M. Berman. The Nucleic Acid Database: new features and capabilities. Nucleic Acids Research, 42(D1):D114–D122, jan 2014. ISSN 0305-1048. doi: 10.1093/nar/gkt980.

22. H. M. Berman, W. K. Olson, D. L. Beveridge, J. Westbrook, A. Gelbin, T. Demeny, S. H. Hsieh, A. R. Srinivasan, and B. Schneider. The nucleic acid database. A comprehensive relational database of three-dimensional structures of nucleic acids. Biophysical Journal, 63(3):751–759, sep 1992. ISSN 00063495. doi: 10.1016/S0006-3495(92)81649-1.

23. Jack A Dunkle, Leyi Wang, Michael B Feldman, Arto Pulk, Vincent B Chen, Gary J Kapral, Jonas Noeske, Jane S Richardson, Scott C Blanchard, and Jamie H Doudna Cate. Structures of the bacterial ribosome in classical and hybrid states of tRNA binding. Science (New York, N.Y.), 332(6032):981–4, may 2011. ISSN 1095-9203. doi: 10.1126/science.1202692.

24. Maria Selmer, Christine M Dunham, Frank V Murphy, Albert Weixlbaumer, Sabine Petry, Ann C Kelley, John R Weir, and V Ramakrishnan. Structure of the 70S ribosome complexed with mRNA and tRNA. Science (New York, N.Y.), 313(5795):1935–42, sep 2006. ISSN 1095-9203. doi: 10.1126/science.1131127.

25. N B Leontis and E Westhof. Geometric nomenclature and classification of RNA base pairs. RNA (New York, N.Y.), 7(4):499–512, apr 2001. ISSN 1355-8382.

26. Wolfram Saenger. Principles of Nucleic Acid Structure. Springer Advanced Texts in Chemistry. Springer New York, New York, NY, 1984. ISBN 978-0-387-90761-1. doi: 10.1007/978-1-4612-5190-3.

27. Saul B. Needleman and Christian D. Wunsch. A general method applicable to the search for similarities in the amino acid sequence of two proteins. Journal of Molecular Biology, 48 (3):443–453, mar 1970. ISSN 0022-2836. doi: 10.1016/0022-2836(70)90057-4.

28. Michiaki Hamada, Hisanori Kiryu, Kengo Sato, Toutai Mituyama, and Kiyoshi Asai. Prediction of RNA secondary structure using generalized centroid estimators. Bioinformatics, 25 (4):465–473, feb 2009. ISSN 1460-2059. doi: 10.1093/bioinformatics/btn601.

29. Bobbie-Jo M. Webb-Robertson, Lee Ann McCue, and Charles E. Lawrence. Measuring Global Credibility with Application to Local Sequence Alignment. PLoS Computational Biology, 4(5):e1000077, may 2008. ISSN 1553-7358. doi: 10.1371/journal.pcbi.1000077.

30. Moeava Tehei, Bruno Franzetti, Dominique Madern, Margaret Ginzburg, Ben Z Ginzburg, Marie-Thérèse Giudici-Orticoni, Mireille Bruschi, and Giuseppe Zaccai. Adaptation to extreme environments: macromolecular dynamics in bacteria compared in vivo by neutron scattering. EMBO reports, 5(1):66–70, jan 2004. ISSN 1469-221X. doi: 10.1038/sj.embor.7400049.

